# A New Species of *Pachytriton* from Anhui, China (Amphibia: Caudata: Salamandridae)

**DOI:** 10.1101/2024.05.08.592934

**Authors:** Zhirong He, Jiaqi Li, Siyu Wu, Shanqing Wang, Na Zhao, Li Ma, Jianping Jiang, Xiaobing Wu, Supen Wang

**Author notes:** These authors contributed equally to this paper (Zhirong He, Jiaqi Li). Author for correspondence. and Brief information of the corresponding author: Xiaobing Wu is a professor at Anhui Normal University. His main research areas include the protection and endangered mechanisms of endangered animals such as Alligator sinensis, population genetic structure and genetic diversity analysis, and biodiversity investigation and monitoring evaluation. Supen Wang is an associate professor at Anhui Normal University. He mainly applies omics techniques to explore the evolutionary mechanisms of terrestrial vertebrates and elucidates the theory of species evolution from a neutral and adaptive perspective, committed to protecting the biodiversity of terrestrial vertebrates. The postal addresses of Professor Wu and Wang are both 241000.

## Abstract

A new species belonging to the genus *Pachytriton* is delineated based on four specimens obtained from the Qingliangfeng Nature Reserve, Huangshan, Anhui, China. The identification of this distinct species within the Southeast Chinese Hilly Area enhances the recognized biodiversity of *Pachytriton* and signifies unresolved interrelations among species in southeastern China. This new species can be distinguished from *P. granulosus* and *P. feii* through meticulous morphological and molecular examinations. The phylogenetic correlation was established utilizing the mitochondrial NADH dehydrogenase subunit 2 gene (ND2). The four recent specimens constitute a cohesive monophyletic clade with robust affirmation. The acknowledgment of this new species elevates the tally of documented *Pachytriton* species to 11.

## 1. Introduction

Strengthening biodiversity conservation and building a community with a shared future for life on earth is a major strategy for China. Amphibians are the most severely threatened group of vertebrates in terms of biodiversity (Luedtke et al., 2023), and the genus *Pachytriton*, as an indispensable part of amphibians, has attracted our attention. The salamandrid genus *Pachytriton* Boulenger, 1878, occurs in eastern and southeastern China, and currently consists of ten nominal species, *P. brevipes* (David, 1875), *P. granulosus* (Chang, 1933), *P. archospotus* (Shen et al., 2008), *P. inexpectatus* (Nishikawa et al., 2011a), *P. feii* (Nishikawa et al., 2011b), *P. moi* (Nishikawa et al., 2011b), *P. changi* (Nishikawa et al., 2012), *P. xanthospilos* (Wu et al., 2012), *P. wuguanfui* (Yuan et al., 2016) and *P. airobranchiatus* (Li et al., 2018). During the global biodiversity crisis, the diversity of amphibians in China has shown a rapid increase. In 2021, 28 new amphibian species (or new records) were added, accounting for 29.47% of the newly added vertebrate species (or new records) in that year (Jiang et al., 2022); In 2022, 44 new amphibian species (or new records) were added, accounting for 38.26% of the newly added vertebrate species (or new records) in that year (Jiang et al., 2023). This highlights the urgent need for comprehensive analysis using morphological and molecular data to further investigate variations within and between species. By exploring the difficult to reach area of Qingliangfeng Nature Reserve, She County, Huangshan City, Anhui province, China and combining the above methods, we have discovered a new species.

We collected a series of *Pachytriton* specimens during the investigation of amphibians in the Qingliangfeng Nature Reserve in the hilly area of southern Anhui. These specimens appear to be very similar to *P. feii* and *P. granulosus* on the surface. Based on the initial description by *P. feii* and *P. granulosus*, as well as our morphological examination and phylogenetic analysis of the collected samples, we believe that these specimens represent a unique unnamed species. Although the number of specimens we have collected is limited and there is no definitive conclusion on the exact location of its population, this study provides a detailed explanation. A complete sequence of the NADH dehydrogenase subunit 2 gene (hereafter, ND2) gene of mitochondrial DNA (mtDNA) of the new species has been deposited in GenBank. Consequently, future studies will be able to detect wild populations of this new species by comparing these GenBank sequences with those of wild specimens collected.

## 2. Materials and methods

### 2.1. Specimens collection

Sampling procedures involving live *Pachytriton* were in accordance with the Wild Animals Protection Law of China and approved by the Animal Ethics Committee at Anhui Normal University. Four unnamed *Pachytriton* specimens were collected from Qingliangfeng Nature Reserve, She County, Huangshan City, Anhui province, China in August 2023. A male adult specimen (collection number: ANU20030001) (Figure 3) and a female adult specimen (collection number: ANU20030002) were obtained on a mountain stream at an altitude of 950 meters in the Qingliangfeng Nature Reserve (Figure 4, C and D). Two adult female specimens (collection numbers ANU20030003, ANU20030004) were obtained from a mountain stream at an altitude of 970 meters in the Qingliangfeng Nature Reserve (Figure 4, A and B, E and F). The specimens were humanely euthanized by immersion in a 95% ethanol solution, then transferred to 75% ethanol for preservation and storage. All these specimens were deposited in Anhui Normal University (ANU), Wuhu, Anhui.

### 2.2. Phylogenetic sampling and analyses

A total of 11 liver samples of the genus *Pachytriton* were used in this study, including four samples of the unnamed specimens from Anhui, three samples of *P. granulosus* from Zhejiang, and four samples of *P. feii* from Anhui (Table 1, Figure 1). All samples were obtained from previously anesthetized and subsequently euthanized specimens and then preserved in 75% ethanol and stored at –80 °C.

**Figure 1.**
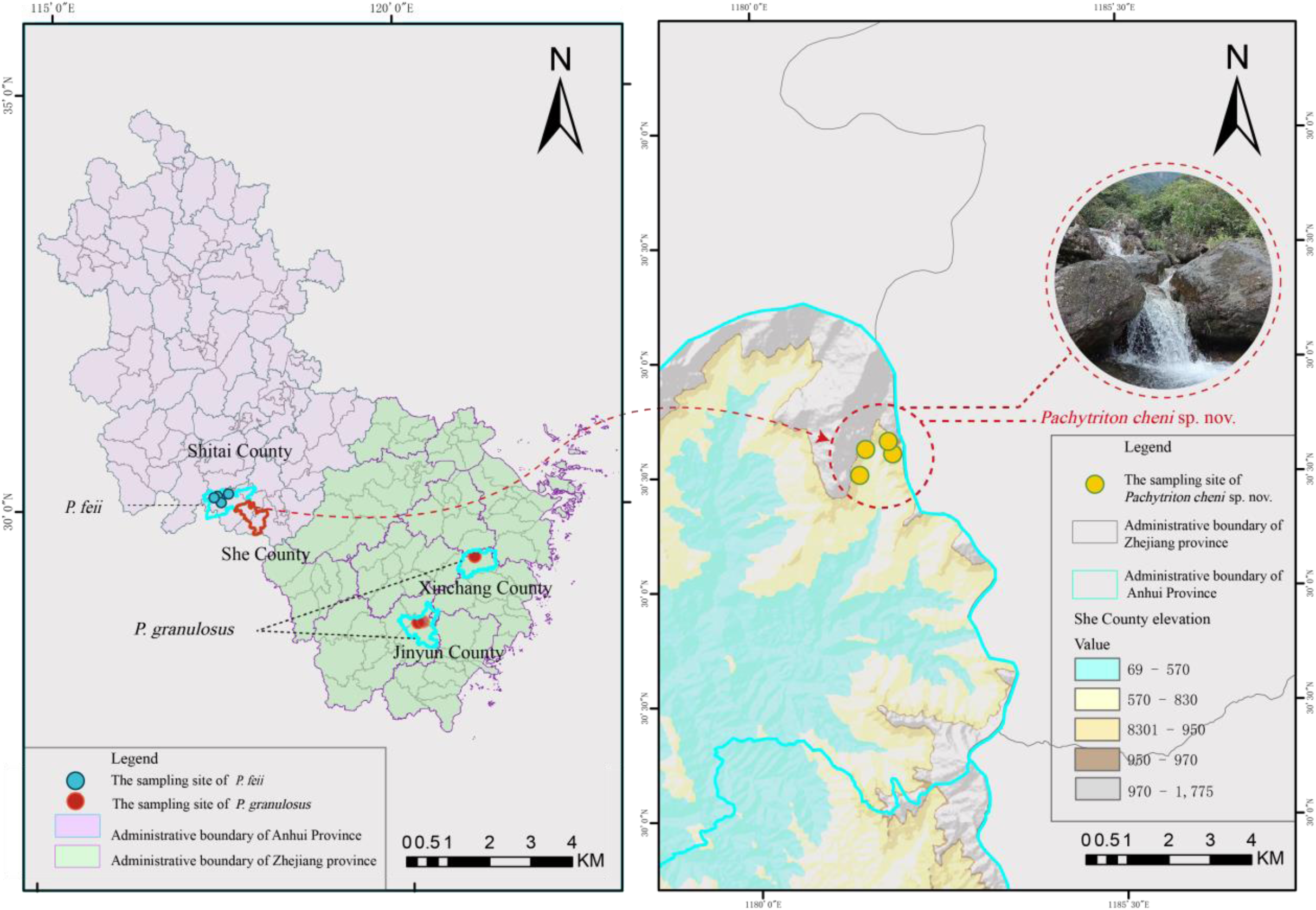
A map of Anhui Province and Zhejiang Province showing sampling localities of *Pachytriton cheni* sp. nov., *P. granulosus* and *P. feii*. A: Maps of Anhui Province and Zhejiang Province. B: Habitat of Qingliangfeng Nature Reserve in She County, Anhui. Solid circles denote sampled localities. For locality information of samples, refer to Table 1.

**Table 1.** Localities, GenBank accession numbers, and Source for all samples used in this study.

| ID no. | Species | Localities | GenBank no. | Source |
| --- | --- | --- | --- | --- |
| 1 | *Pachytriton cheni* sp. nov. | China: Anhui: Huangshan: She | SRR28764900 | This study |
| 2 | *Pachytriton cheni* sp. nov. | China: Anhui: Huangshan: She | SRR28764897 | This study |
| 3 | *Pachytriton cheni* sp. nov. | China: Anhui: Huangshan: She | SRR28764899 | This study |
| 4 | *Pachytriton cheni* sp. nov. | China: Anhui: Huangshan: She | SRR28764896 | This study |
| 5 | *Pachytriton feii* | China: Anhui: Chizhou: Shitai | SRR28764895 | This study |
| 6 | *Pachytriton feii* | China: Anhui: Chizhou: Shitai | SRR28764894 | This study |
| 7 | *Pachytriton feii* | China: Anhui: Chizhou: Shitai | SRR28764893 | This study |
| 8 | *Pachytriton feii* | China: Anhui: Chizhou: Shitai | SRR28764892 | This study |
| 9 | *Pachytriton granulosus* | China: Zhejiang: Shaoxing: Xingchang | SRR28764891 | This study |
| 10 | *Pachytriton granulosus* | China: Zhejiang: Lishui: Jinyun | SRR28764890 | This study |
| 11 | *Pachytriton granulosus* | China: Zhejiang: Lishui: Jinyun | SRR28764898 | This study |
| 12 | *Pachytriton archospotus* | China: Jiangxi: Pingxiang: Wugongshan | JX907837 | Wu et al. (2013) |
| 13 | *Pachytriton archospotus* | China: Jiangxi: Pingxiang: Wugongshan | JX907838 | Wu et al. (2013) |
| 14 | *Pachytriton archospotus* | China: Jiangxi: Pingxiang: Wugongshan | JX907839 | Wu et al. (2013) |
| 15 | *Pachytriton archospotus* | China: Jiangxi: Jinggangshan: Jinggangshan | JX907840 | Wu et al. (2013) |
| 16 | *Pachytriton archospotus* | China: Jiangxi: Jinggangshan: Jinggangshan | JX907841 | Wu et al. (2013) |
| 17 | *Pachytriton archospotus* | China: Jiangxi: Jinggangshan: Jinggangshan | JX907842 | Wu et al. (2013) |
| 18 | *Pachytriton archospotus* | China: Hunan: Guidong: Qiyunshan | GQ303628 | Wu et al. (2010) |
| 19 | *Pachytriton archospotus* | China: Hunan: Guidong: Qiyunshan | GQ303629 | Wu et al. (2010) |
| 20 | *Pachytriton archospotus* | China: Hunan: Guidong: Qiyunshan | GQ303630 | Wu et al. (2010) |
| 21 | *Pachytriton archospotus* | China: Guangdong: Shixing: Guanyindong | JX237731 | Wu et al. (2012) |
| 22 | *Pachytriton archospotus* | China: Hunan: Guidong: Bamianshan | KT152558 | Wu et al. (2015) |
| 23 | *Pachytriton archospotus* | China: Jiangxi: Pingxiang: Wugongshan | KT152559 | Wu et al. (2015) |
| 24 | *Pachytriton archospotus* | China: Jiangxi: Jinggangshan: Jinggangshan | KT152560 | Wu et al. (2015) |
| 25 | *Pachytriton moi* | China: Guangxi: Guilin: Xinan; Mt. Mao'er | JX237746 | Wu et al. (2012) |
| 26 | *Pachytriton moi* | China: Guangxi: Guilin: Xinan; Mt. Mao'er | JX237747 | Wu et al. (2012) |
| 27 | *Pachytriton moi* | China: Guangxi: Guilin: Xinan; Mt. Mao'er | KT152556 | Wu et al. (2015) |
| 28 | *Cynops ensicauda* | Japan | DQ517769 | Weisrock et al. (2006) |
| 29 | *Echinotriton andersoni* | Japan | DQ517774 | Weisrock et al. (2006) |

Genomic DNA was extracted from frozen or ethanol-preserved tissues using standard proteinase K extraction procedures. One mitochondrion gene, namely NADH dehydrogenase subunit 2 (ND2), was amplified using the primers ND2-4F (5′-TATGAGTACGAGCATCATACCC-3′) and ND2-4R (5′-CTTCTGCTTAAGACTTTGAAGGTC-3′) (Lyu et al., 2021). PCR amplifications were processed with the following cycling conditions: denaturing at 95 °C for 4 min, 35 cycles of denaturing at 95 °C for 40 s, annealing at 53 °C for 34 s and extending at 72 °C for 60 s, and a final extending step at 72 °C for 10 min. The PCR products were purified using a rotating column and then sequenced using forward and reverse primers on the ABI Prism 3730XL automated DNA sequencer by Anhui General Biology Co., Ltd. All obtained sequences have been stored in GenBank (Table 1).

Thirteen sequences of *P. archospotus*, three sequences of *P. moi* and two sequences of out-group species from the genera *Pachytriton* Boulenger, 1878, were retrieved from GenBank and included in our dataset for phylogenetic analyses. Detailed information is provided in Table 1. The DNA sequences were aligned using the Clustal W algorithm with default parameters (Thompson et al., 1997). The most suitable model for nucleotide evolution for each gene’s codon position and tRNA was determined using jModelTest 2.1.7 (Darriba et al., 2012), based on the Akaike Information Criterion. The optimal model selected for each partition was the GTR+I model. The sequenced data were subjected to maximum likelihood (ML) analysis in MAGA11. In the ML analysis, a bootstrap consensus tree generated from 1000 replicates was utilized to depict the evolutionary lineage of the analyzed taxa. We regarded tree topologies with bootstrap values (BS) of 70% or greater as sufficiently supported (Huelsenbeck and Hillis, 1993).

### 2.3 Morphological comparison and analyses

In morphological comparative analysis, the selected target traits can be roughly divided into two types: quantitative traits and qualitative traits. Quantitative traits refer to the continuous distribution of variation in a population of a certain trait, which cannot be clearly grouped. Quality traits refer to traits that can be qualitatively identified and clearly grouped within a population, with significant differences between groups.

#### Quantitative analysis

For the analysis of quantitative traits, there are four unnamed specimens, four *P. feii* and three *P. granulosus* were measured externally using a digital caliper (Deli DL91200 stainless steel 200mm Digita vernier caliper) accurate to 0.01 millimeters. Only adult specimens were measured. These measurements are as follows: total length (TOL) from tip of snout to tip of tail; snout–vent length (SVL) from tip of snout to posterior edge of vent; tail length (TAL) from posterior edge of vent to tip of tail; maximum tail depth (TAD); head length (HL) from tip of snout to the posterior edge of the parotoid gland; maximum head width (HW); snout length (SL) from tip of snout to the anterior corner of eye; eye diameter (ED) from the anterior corner to the posterior corner of the eye; interorbital distance (IOD) between the anterior corner of each eye; eye–nostril distance (END) from the anterior corner of the eye to the nostril; internasal distance (IND) between the external nares; axilla–groin length (AG) between the axilla and the groin along the body; forelimb length (FLL) from elbow to tip of finger III; and hindlimb length (HLL) from knee to tip of toe III. Symmetric head features are only measured on the right side, while asymmetric features are recorded on both sides and averaged. Statistical analyses on the morphometric measurements were performed in R version 4.3.2 (Lyu et al., 2020). Due to evident sexual size dimorphism (Fei et al., 2006), separate analyses were performed for males and females. All measurements were lntransformed to normalize and reduce the variance. One-way analysis of variance (ANOVA) was conducted with statistically similar variances (*P* values > 0.05 in the Levene’s test) using car R package. Principal component analysis (PCA) was performed to reduce the dimensionality of variation in the data to find whether morphological variation forms the basis of detectable group structure, using prcomp function and ggplot2 package.

#### Qualitative analysis

We used the 14 morphological features mentioned above to rate each individual. When making comparisons, we considered only those characters that were present in all members of the population and dismissed those with variation among individuals, in order to describe discrete and diagnostic characters (following Wiens 2000 criteria).

## 3. Results

### 3.1 Molecular phylogenetic relationships

Molecular analysis was performed on the mitochondrial ND2 gene fragments of thirteen known *P. archospotus*, three known *P. moi* and two known out-group species *Cynops ensicauda* and *Echinotriton andersoni*, as well as four unnamed specimens, four *P. feii* and three *P. granulosus* in this study. A phylogenetic tree was constructed based on the maximum likelihood method (ML), and the obtained topological structure is shown in Figure 2. The topology structure obtained from ML analysis is shown in Figure 2. The P-distances based on the ND2 gene among species are presented in Table 2. As shown in the tree, three clades (BS 90) were revealed for the samples of *Pachytriton*. The first clade consists of *P. granulosus*, unnamed samples from Qingliangfeng Nature Reserve, and *P. moi*, but with relatively weak support values (BS 60). The unnamed samples from the Qingliangfeng Nature Reserve formed a unique lineage (BS 96) that is relatively close to *P. granulosus* from Zhejiang, China. The second clade is composed of *P. archospotus* (BS 99). The third branch is composed of *P. feii* (BS 100).

**Figure 2.**
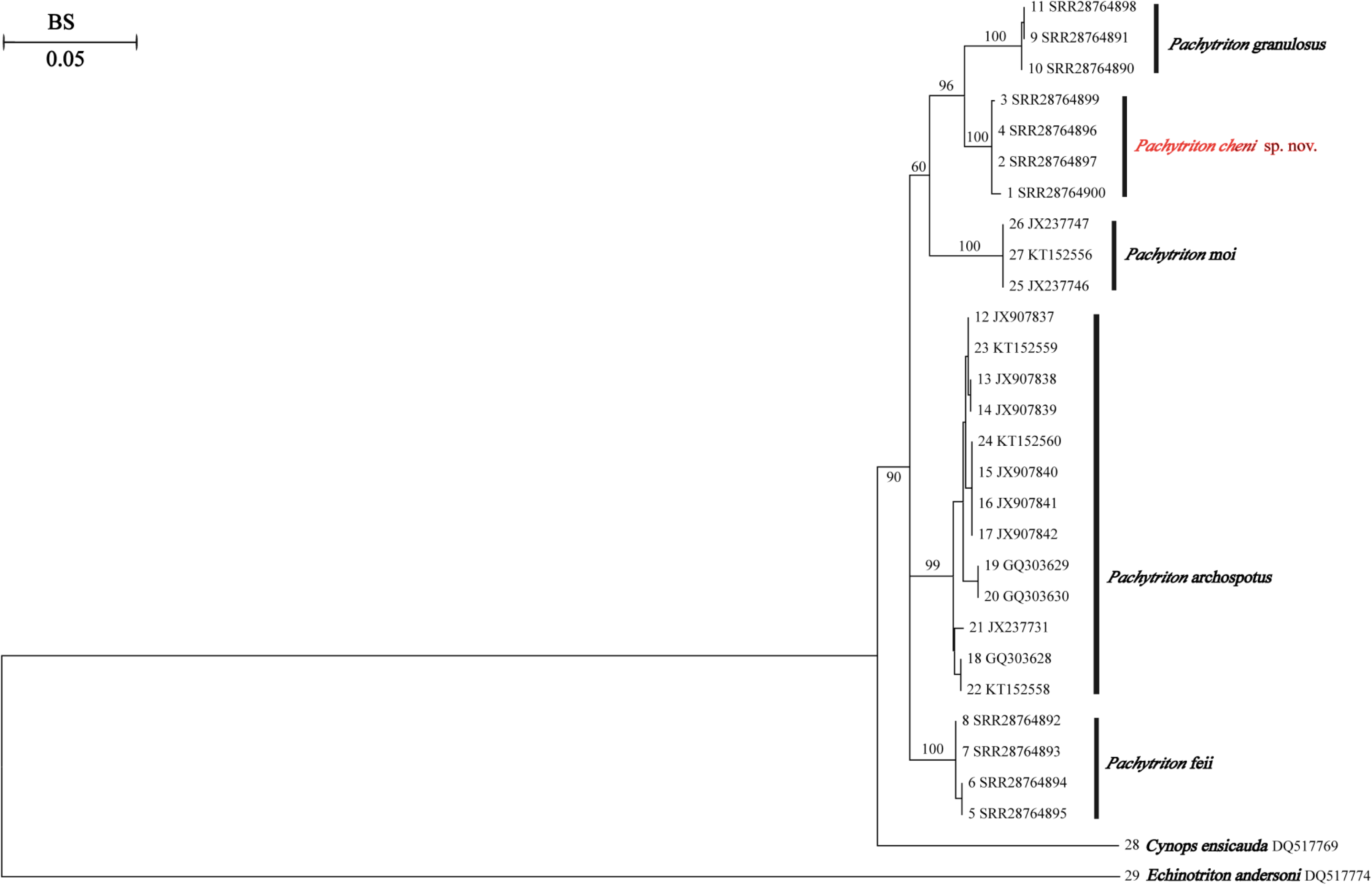
Maximum-likelihood phylogenies based on mitochondrial ND2 gene. Bootstrap supports (BS) are all shown. ID number in Table 1 and GenBank number are displayed at the end of the lineages.

**Table 2.** Mean *P*-distance based on the ND2 gene among *Pachytriton* species (in 1%).

| ID | Species | 1 | 2 | 3 | 4 | 5 |
| --- | --- | --- | --- | --- | --- | --- |
| 1 | *Pachytriton moi* | — |  |  |  |  |
| 2 | *Pachytriton archospotus* | 5.13 |  |  |  |  |
| 3 | *Pachytriton cheni* sp. nov. | 5.07 | 5.15 |  |  |  |
| 4 | *Pachytriton feii* | 5.39 | 4.00 | 4.50 |  |  |
| 5 | *Pachytriton granulosus* | 5.50 | 6.47 | 3.46 | 5.38 | — |

### 3.2 Morphological comparisons

Morphologically, the *Pachytriton* specimens from these four unnamed lineages can be distinguished from all known congeners reliably (details in the Taxonomic account below). The unnamed *Pachytriton* population from She County, Huangshan City and the sister taxon *P. feii* from She County, Chizhou City and *P. granolosus* from Zhejiang Province were statistically analyzed by morphometric measurements (Table 3; Figure 3). Since only one male specimen of the unnamed *Pachytriton* population, one male specimen of *P. feii,* and no male specimens of *P. granulosus* were collected, related morphometric analyses on males were not carried out.

**Figure 3.**
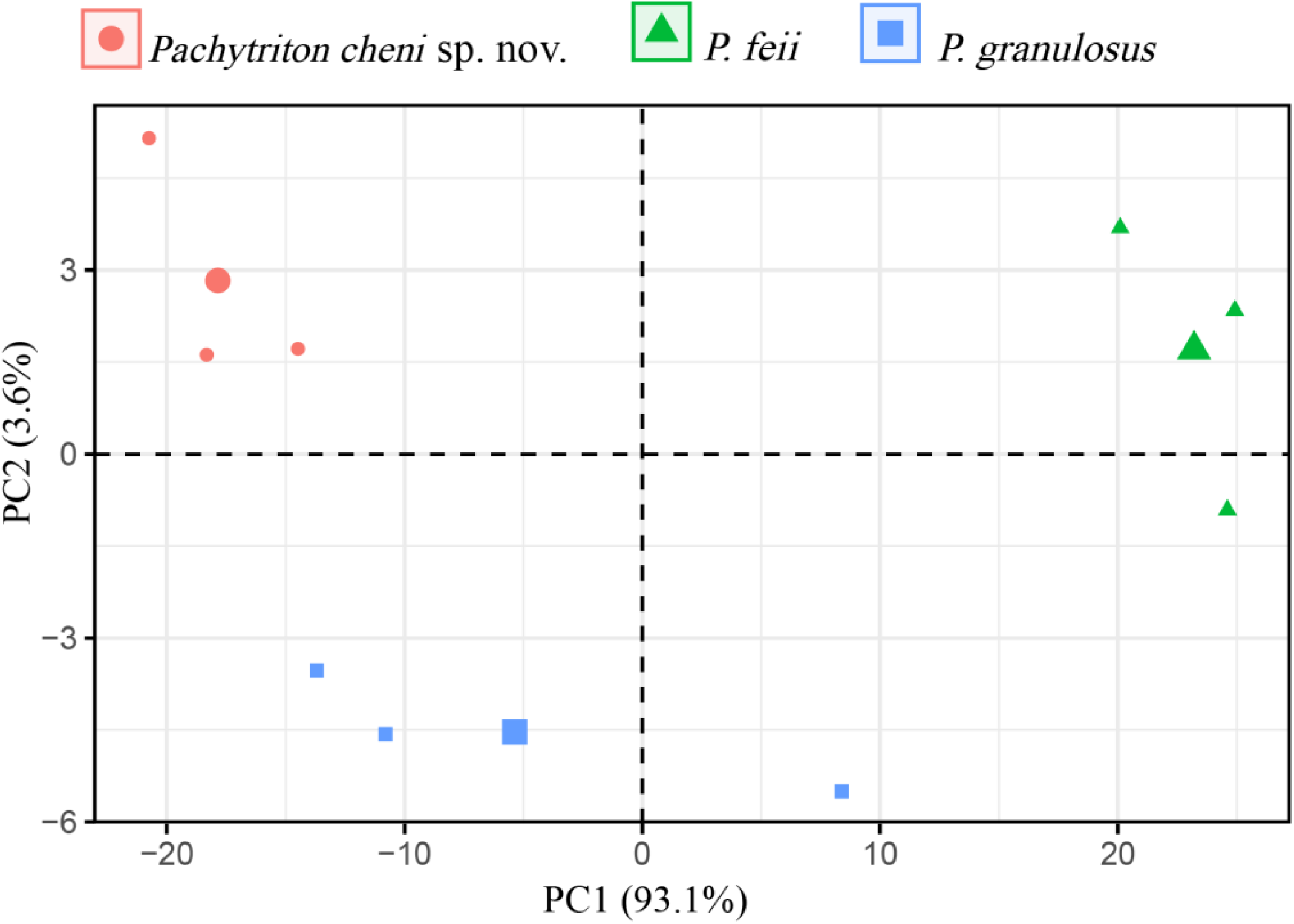
Scatter plot of PC1 and PC2 of Principal Component Analysis based on the morphometric measurements, distinguishing *Pachytriton cheni* sp. nov. and *P. feii*.

**Table 3.** Morphometric comparisons based on the morphometric measurements (in mm) of *Pachytriton cheni* sp. nov., *P. feii* and *P. granulosus.* The P-value refers to the P-value of *Pachytriton cheni* sp. nov. and *P. Feii* and the P-value’ refers to the P-value of *Pachytriton cheni* sp. nov. and *P. granulosus.* * *P*-values < 0.05, ** *P* - values < 0.01, *** *P* -values < 0.001.

| Species | *Pachytriton cheni* sp. nov. |  | *P. feii* |  | *P*-values | *P. granulosus* | *P*-values' |
| --- | --- | --- | --- | --- | --- | --- | --- |
| Gender | Male(n=1) | Female(n=3) | Male(n=1) | Female(n=3) | Female | Female(n=3) | Female |
| TOL | 120.58 | 131.23-149.48(138.25±96.52) | 164.25 | 156.18-161.23(158.84±6.43) | 0.0246* | 121.75-130.86(126.75±21.35) | 0.14 |
| SVL | 65.17 | 71.89-76.64(73.67±6.70) | 90.15 | 85.43-88.42(86.88±2.24) | 0.00157** | 68.9-69(68.97±0) | 0.0347* |
| TAL | 55.41 | 60.51-73.24(65.57±45.60) | 74.10 | 73.44-77.81(76.04±5.29) | 0.0639 | 58.65-62.75(61.12±4.73) | 0.338 |
| TAD | 8.16 | 7.20-9.94(8.39±1.97) | 10.56 | 7.80-10.75(9.23±2.18) | 0.515 | 10.85-12.07(11.37±0.4) | 0.0286* |
| HL | 19.42 | 18.67-20.71(19.86±1.13) | 23.25 | 20.76-22.88(22.13±1.41) | 0.0692 | 17.46-19.88(18.56±1.5) | 0.238 |
| HW | 11.88 | 11.73-12.70(12.10±0.27) | 15.49 | 13.81-15.49(14.75±0.74) | 0.0103* | 12.03-13.11(12.72±0.36) | 0.249 |
| SL | 4.30 | 3.84-4.35(4.11±0.07) | 5.95 | 5.26-5.95(5.56±0.12) | 0.00446** | 5.57-5.87(5.7±0.02) | 0.000772*** |
| ED | 3.18 | 2.62-3.67(3.13±0.28) | 4.30 | 3.05-4.05(3.57±0.25) | 0.353 | 3.71-4.37(4.1±0.12) | 0.0557 |
| IOD | 6.94 | 6.46-7.71(7.27±0.50) | 5.75 | 5.40-6.59(5.82±0.45) | 0.0605 | 4.45-4.76(4.6±0.02) | 0.00305** |
| EN | 3.77 | 3.47-4.33(3.90±0.18) | 5.74 | 3.80-4.46(4.06±0.12) | 0.63 | 4.55-4.84(4.68±0.02) | 0.04* |
| IND | 3.58 | 4.84-5.16(4.97±0.03) | 4.31 | 4.24-4.86(4.49±0.11) | 0.0861 | 3.71-4.14(3.97±0.05) | 0.00354** |
| AG | 29.81 | 33.58-38.26(35.27±6.73) | 46.15 | 43.98-48.13(46.29±4.47) | 0.00468** | 35.67-38.67(36.69±2.93) | 0.473 |
| FLL | 9.85 | 9.89-10.75(10.35±0.19) | 14.15 | 11.20-13.03(12.22±0.87) | 0.0349* | 12.21-14.62(13.28±1.51) | 0.0176* |
| HLL | 12.72 | 11.50-12.40(12.03±0.23) | 16.34 | 13.68-15.42(14.64±0.79) | 0.0109* | 13.81-15.62(14.63±0.84) | 0.0121* |

The results of ANOVA on morphometrics for females showed that individuals of the unnamed *Pachytriton* population and *P. feii* are significantly different in TOL, SVL, HW, AG, FLL and HLL for females (*P*-values < 0.05), and significant differences in SVL, TAD, SL, IOD, EN, IND, FLL and HLL between the unnamed *Pachyton* population and *P. granulosus* (P-values < 0.05). For the PCA result of females, the first two principal components (PCs) account for 93.1% and 3.6% of the variance, respectively (in total > 90.0% of the variance). As illustrated in the scatter plots (Figure 3), the female individuals of the unnamed *Pachytriton* population, *P. feii* and *P. granulosus* formed respective clusters and were clearly separated.

Combined with the results above morphological examination, morphometric statistical analysis and phylogenetic analysis, this *Pachytriton* population from She County, Huangshan City is considered as a new species described in this paper.

## 4. Taxonomic accounts

### Holotype

ANU 20230001 (Figure 4), an adult male, collected by Zhirong He and Siyu Wu on 24 August 2023 at a mountain stream at an altitude of 950 meters in Qingliangfeng Nature Reserve (30.08445491°N, 118.86914777°E), She County, Huangshan City, Anhui province, China. When collecting, the depth of the mountain stream water flow is shallow, but the flow velocity is fast. The GenBank accession number of ND2 sequence is SRR28764900.

**Figure 4.**
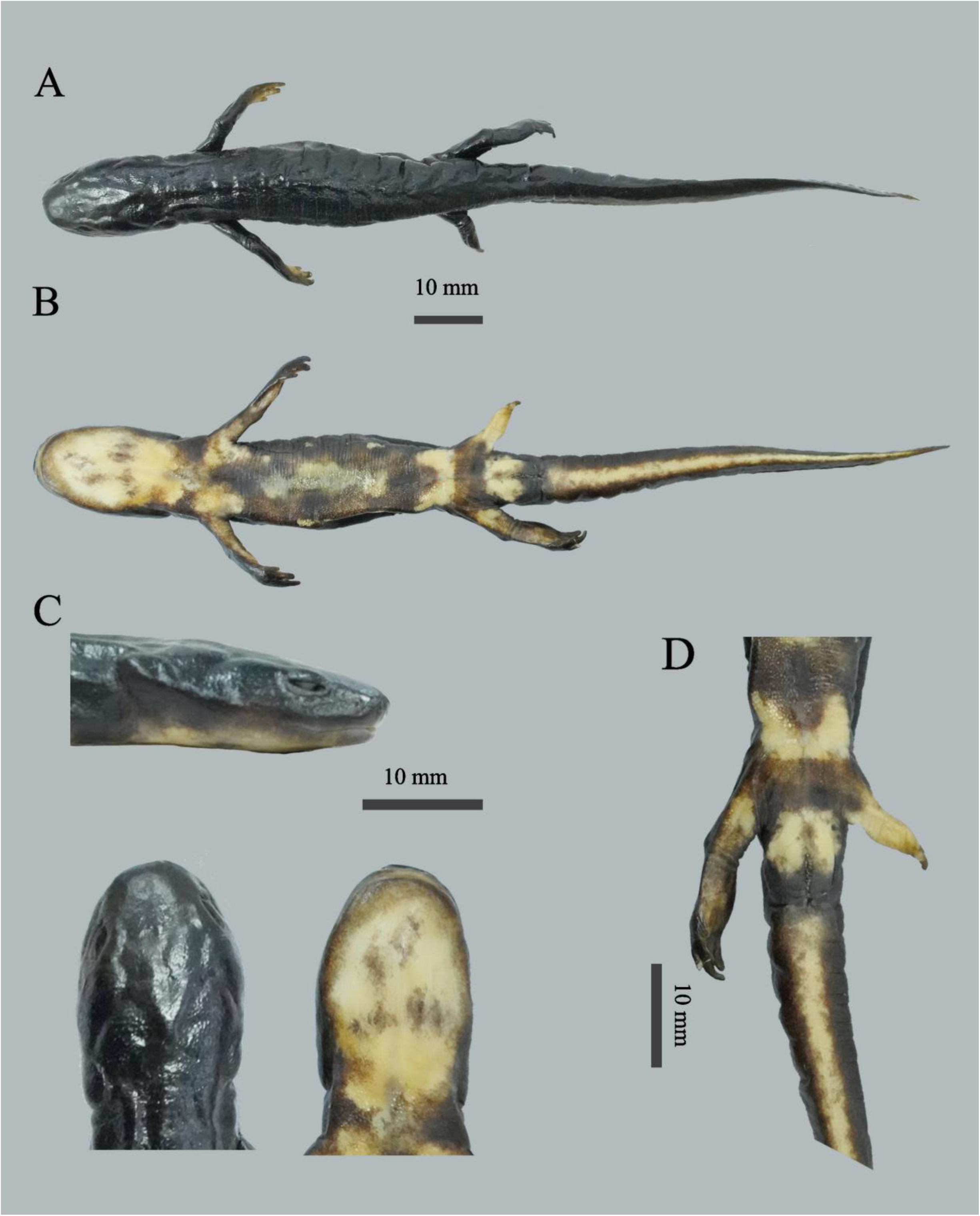
ANU 20230001 of *Pachytriton cheni* sp. nov. in preservative. A: dorsolateral view. B: ventral view. C: lateral, dorsal, and ventral views of head. D: region around cloaca. Photos by SY Wu.

### Paratype

ANU 20230002 (Figure 5, A and B), an adult female, collected by Zhirong He and Siyu Wu on 24 August 2023 at a mountain stream at an altitude of 970 meters in Qingliangfeng Nature Reserve (30.08494572°N, 118.86950445°E), She County, Huangshan City, Anhui province, China. The surrounding environment of the collection point is a quiet pond in a mountain stream, with deep water and many stones at the bottom. The GenBank accession number of ND2 sequence is SRR28764897.

**Figure 5.**
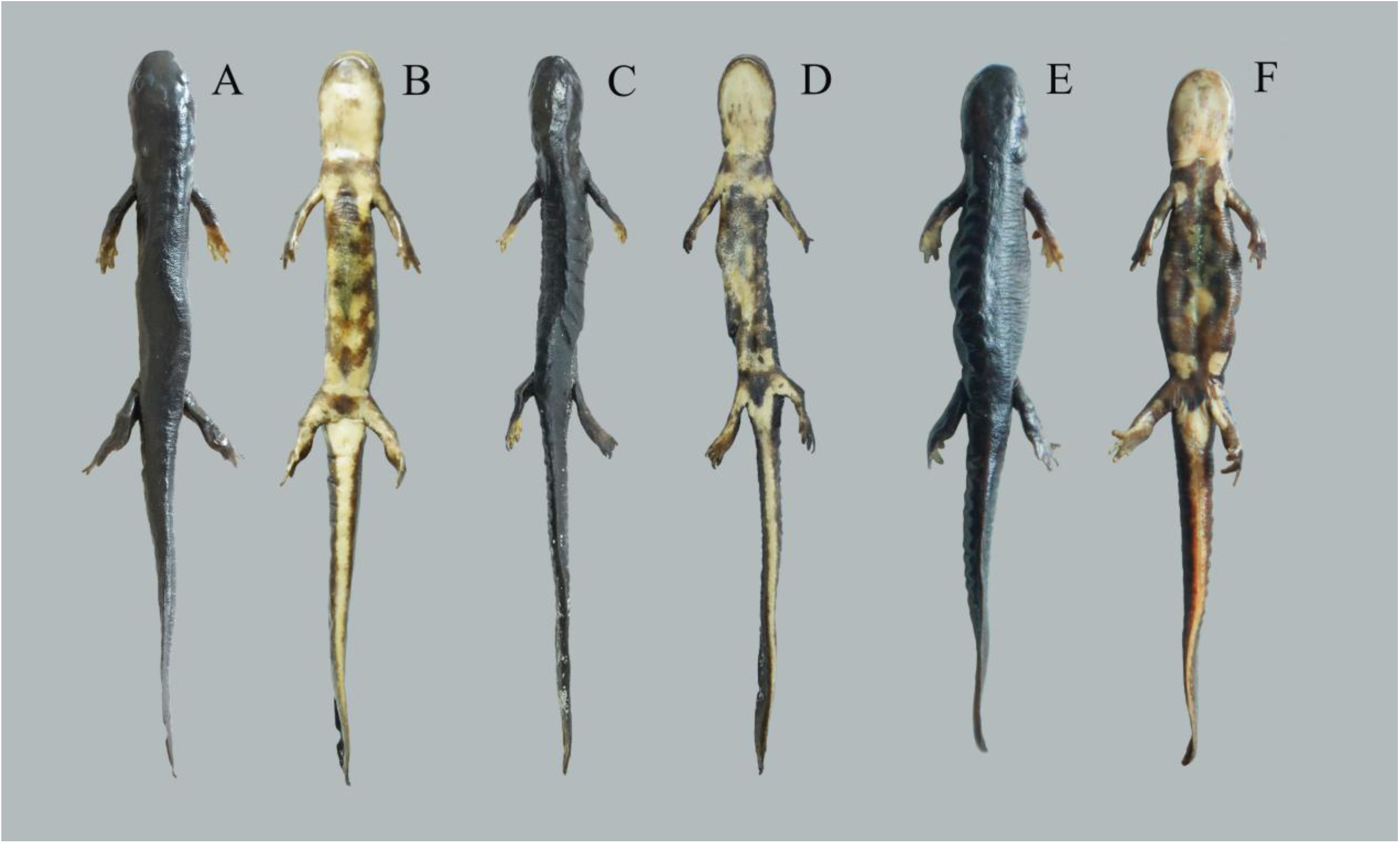
ANU 20230002(A and B), ANU 20230003 (C and D) and ANU 20230004 (E and F) of *Pachytriton cheni* sp. nov. in preservative. Dorsolateral views (A, C and E) and ventral views (B, D and F). Photos by SY Wu. Scale bar=10 mm.

ANU 20230003 (Figure 5, C and D), an adult female, same collection data as the holotype. The GenBank accession number of ND2 sequence is SRR28764899.

ANU 20230004 (Figure 5, E and F), an adult female, same collection data as the specimen ANU 20230002. The GenBank accession number of ND2 sequence is SRR28764896.

### Etymology

Professor Bi-Hui Chen, a world-renowned herpetologist, devoted his whole life to the education and scientific research of vertebrates. He is known as the "father of the Chinese alligator" because he obtained the survival strategy of the critically endangered species of *Alligator sinensis* from nature through personal practice, and ultimately solved the endangered problem of *Alligator sinensis*. He and his colleagues published "Anhui Amphibians and Reptiles Fauna" (Chen, 1991), which made important contributions to the research of amphibians and reptiles in Anhui Province and China. To commemorate Professor Bi-Hui Chen, who passed away in 2022, we suggest naming the unnamed *Pachytriton* species discovered this time after the surname of Professor Bi-Hui Chen, which is commonly known in English as "*Pachyton cheni*" and in Chinese as “陈氏肥螈( chén shì féi yuán)”.

### Diagnosis

(1) a small-sized newt of the genus *Pachytriton*, TOL is 120.58 mm in the adult males, TOL is 131.23-149.48 mm in the adult females; (2) body slender, slim limbs, slightly flattened back and abdomen; (3) head oval and narrow, head length greater than head width; (4) short snout, blunt and round snout; (5) surface smooth, gular fold absent; (6) the nostrils are extremely close to the snout end; (7) well-developed labial folds; (8) upper and lower jaw with fine teeth; (9) significant neck folds; (10) both the forelimbs and hindlimbs are relatively short; (11) finger formula: 3>2>4>1; Toe style: 3>4>2>5>1; (12) the base of the tail is wide and thick, the latter half gradually flattens to the side, and the end is blunt and round; (13) dorsum, flanks, limbs, and upper side of tail usually uniformly dark; (14) ground color of venter dark brown, with irregular bright orange patches on chin, chest, and belly.

### Description of the holotype

ANU 20230001 (Figure 4), adult male with a slender and small-sized body (TOL 120.58 mm, SVL 65.17 mm, TAL 56.76 mm). Skin smooth; head oval in dorsal view and nearly flat in profile; hnout truncate, protruding slightly beyond lower jaw; nostrils close to snout tip; parotoid region evident; cloacal opening oval, slightly protruding; tail laterally compressed and the end is blunt and round.

### Coloration of the holotype

In life, dorsum, flanks, limbs, and upper side of tail uniformly black; vertebral ridge uniformly black as dorsal surface; ground color of venter dark brown, with irregular bright orange patches on chin, chest, and belly; cloacal opening and underside of tail bright orange. In preservative (Figure 4), dorsum, flanks, and limbs uniformly dark. Ventral bright markings fading to cream.

### Variations

Measurements of the type series are given in Table 3. Females (TOL 131.23-149.48 mm) distinctly larger than males (TOL 120.58 mm). The cloaca is wider and more swollen in males than in females; irregular bright orange patches on ventral surface vary among individuals. Compared with the holotype, all paratype have fewer bright orange spots around the chin, abdomen, and cloaca; but in other forms, it is very similar to the holotype.

### Distribution and habitat

All individuals were observed in a mountain stream on a peak at late August. *Pachytriton cheni* sp. nov. is currently known only from its type locality in Qingliangfeng Nature Reserve in southern Anhui(Figure 6). This new species inhabits small montane streams (1–2 m wide) in broadleaf forests near the top of the mountain at elevations ranging from 900 m to 1000 m. Large boulders are scattered throughout the stream. The flowing water is very clear, and some areas have formed small ponds after the rain. Water temperatures ranged from 17.5 °C to 18.9 °C in the night of 24 August 2023. The pH value of water was close to 7.6 according to PH meter. Stream substrates include gravels, scattered small rocks, leaves and sands. The surrounding forests are mainly composed of *Emmenopterys henryi Oliv.*, *Alnus rubra*, *Pseudolarix amabilis (J. Nelson) Rehder*, *Quercus stewardii Rehder* and *Phyllostachys edulis (Carrière) J. Houz*. Other amphibians and reptiles that co-inhabit the stream include *Amolops wuyiensis, Quasipaa exilispinosa, Trimeresurus stejnegeri* and *Deinagkistrodon acutus*.

**Figure 6.** Habiat at the *Pachytriton cheni* sp. nov. in Qingliangfeng Nature Reserve, Huangshan, Anhui, China.

### Conservation recommendation

Due to the Qingliangfeng Nature Reserve being located in the southern part of China, the *Pachytriton* genus is generally suitable for cold water (Wu et al. 2013), high annual temperature restricts the distribution of the new species to high elevations near the top of the mountain. Individuals of this newt sometimes gather together in deeper pools, but its population size seems much smaller compared to other more common species such as *P. feii* and *P. granulosus*. Even though the type locality of *Pachytriton cheni* sp. nov. is well protected by the Qingliangfeng Nature Reserve and no major threat factors have been observed, the extent of occurrence of this species estimated to be less than 100 km2, and the area of occupancy is estimated to be less than 10 km^2^. This new species belongs to *Pachytriton* in amphibians, which has been facing severe survival pressure recently. Luedtke et al. (2023) integrated data from GAA1 and the Second Global Amphibian Assessment (GAA2) in 2022 and pointed out in the journal *Nature* that the pathogen of chytrid is gradually becoming a new threat to amphibians. This new species may be infected with *Batrachochytrium dendrobatidis* and *B. salamandrivorans*, which will pose a great threat to its population size (Wang et al., 2017; He et al., 2024). Therefore, we suggest that the exact locality of the new species should be determined as soon as possible and relevant pathogen screening should be conducted.

## 5. Discussion

*Pachytriton* often lives in mountain streams. *P. feii* and *P. granulosus* are geographically distributed in Huangshan City, Anhui Province (*P. feii*: Frost, 2022; *P. granulosus*: Fei et al., 2012). The mountain streams in the hilly areas of southeast Chinese are rarely visited by people, and there may be potential species different from the *P. feii* and *P. granulosus*. Therefore, further research is needed on the phylogenetic relationship and population division between the genus *Pachytriton* in the Southeast Chinese Hilly Area.

The Qingliangfeng Nature Reserve is located at the border of Anhui and Zhejiang, and belongs to the southeast hilly area. It has an ancient history, complex landforms, and abundant mountain and stream resources, which is very suitable for the living environment of *Pachytriton*. Four specimens (ANU20230001、ANU20230002、ANU20230003、ANU20230004) sampled from Qingliangfeng Nature Reserve were also used in detailed morphological and molecular studies of Chinese amphibian.

The morphological analysis results revealed significant differences between the new species and *P. feii* and *P. granulosus* (Table 4). *Pachytriton cheni* sp. nov. is phylogenetically closely related to *P. granulosus,* which is distributed in Zhejiang; however, it possesses the following distinct characteristics: a longer total body length; a more slender body shape; a head length significantly greater than its width; a shorter snout; nostrils positioned at the anterior end of the snout; relatively short forelimbs and hindlimbs, yet not stocky; a dark brown coloration on the dorsal side, flanks, and tail of the body; and rare orange-red spots on the back. *Pachytriton cheni* sp. nov. can be readily distinguished from *P. feii* by the following morphological features: a shorter total body length; a more slender body shape; a shorter and sharper snout; and an inconspicuous V-shaped protrusion. Moreover, in contrast to the light orange-red or orange-yellow spots on the ventral side of *P. feii*, the ventral side of *Pachytriton cheni* sp. nov. exhibits orange-red coloration with some brown short lines or worm-like spots.

**Table 4.**
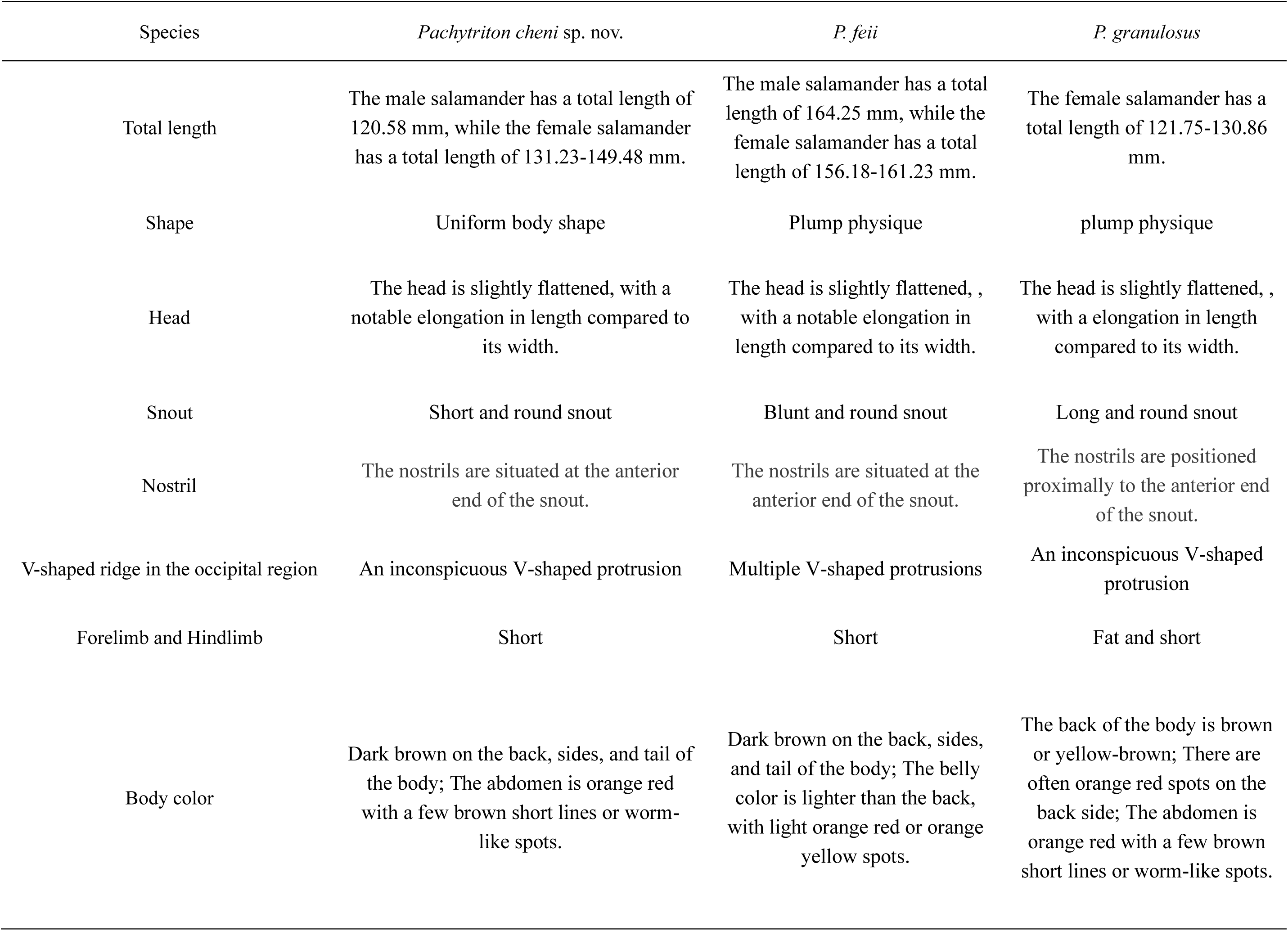
Comparison and presentation of the main quality characteristics of *Pachytriton cheni* sp. nov., *P. feii* and *P. granulosus*.

The phylogenetic analysis results indicate that the specimens discovered in the Qingliangfeng Nature Reserve have formed a unique lineage (BS 96) and are closely related to *P. granulosus* in Zhejiang. Although the bparanching relationship between these four specimens and other species of the genus *Pachytriton* has not been resolved, the formation of independent branches between different species is beyond doubt. So, the specimens from Qingliangfeng Nature Reserve should be a potential new species of *Pachytriton*.

At the species level, “subdivision” is recommendable. This approach aids in a more effective and precise understanding and description of the natural history of species, as well as facilitating consistent communication and actions in taxonomy and conservation biology practices (Huang et al., 2021). Despite the incompleteness of morphological analysis due to insufficient male specimens, considering phylogenetic and morphological differences, it is plausible to regard the unnamed population as an independent species. With the inclusion of the newly described species in this study, the genus *Pachytriton* now comprises 11 recognized species.

Geodiversity is the foundation of biodiversity (Ren et al., 2021). The southeastern hilly area, the largest and most densely populated hilly region in the country, China, possesses the largest land area. The Huangshan area has even been suggested as a Pleistocene refugium (Wu et al., 2013). The Qingliangfeng Nature Reserve, which is rarely visited, contains abundant biodiversity (Weng et al., 2009; Guo et al., 2016), including diverse lineages and high genetic diversity of various organisms. This study further highlights the role of reserve in promoting regional genetics and biodiversity. Therefore, we advocate for increased efforts to protect the Qingliangfeng Nature Reserve and determine the habitat of the newly discovered species *Pachytriton cheni* sp. nov. to facilitate further research on its systematic evolution and biogeography.

## Acknowledgements

This research was funded by the National Natural Science Foundation of China (31901120 and 31700320), Beijing Natural Science Foundation (5192016), Anhui Provincial Key Laboratory of the Conservation and Exploitation of Biological Resources (swzy202006). We thank anonymous reviewers for their valuable suggestions on this work.

## References

AmphibiaWeb. 2022. AmphibiaWeb: information on amphibian biology and conservation. Berkeley, USA: University of California. Available from: http://amphibiaweb.org/ (accessed 10 October 2022)

Chang M. L. Y. 1933. On the salamanders of Chekiang. Contri Biol Lab Sci Soc China, Nanjing 9: 305–328.

Chen B. H. 1991. Anhui Amphibians and Reptiles Fauna. Hefei: Anhui Science and Technology Press. (in Chinese)

Darriba D., Taboada G. L., Doallo R. & Posada D. 2012. jModelTest 2: more models, new heuristics and parallel computing. Nat. Methods 9: 772.

David A. 1875. Journal de mon Troisième Voyage d’Exploration dans l’Empire Chinoise. Vol. 1. Hachette, Paris, 383 pp.

Fei L., Hu S. Q., Ye C. Y., Huang Y. Z. 2006. Fauna Sinica. Amphibia. Volume 1. General Accounts of Gymnophiona and Urodela. Beijing, China: Science Press.

Fei L., Ye C. Y., Jiang J. P. 2012. Colored Atlas of Chinese Amphibians and Their Distributions. Chengdu: Sichuan Scientific & Technical Publishers, 436–471.

Frost D. R. 2022. Amphibian Species of the World 6.1, an Online Reference. New York, USA: American Museum of Natural History. Availabl from: https://amphibiansoftheworld.amnh.org/ (accessed 10 October 2022)

Guo R., Wang Y., Weng D., Cheng Z., Wang J., & Wang X. 2016. *Carabid beetle* (Coleoptera, Carabidae) species diversity and environmental factors in biotopes of Zhejiang Qingliangfeng National Nature Reserve, China. Journal of Zhejiang A&F University, 33(4), 551–557.

He Z. R., Wu S. Y., Shi Y. Y., Wang Y. T., Jiang Y. X., Zhang C. N., Zhao N., Wang S. P. 2024. Current status and challenges on the effects of chytrid infection on amphibian populations. Biodiversity Science, 32, 23274.

Huang R., Peng L., Yu L., Huang T. Q., Jiang K., Ding L., Chang J. K., Yang D. C., Xu Y. H., Huang S. 2021. A new species of the genus *Achalinus* from Huangshan, Anhui, China (Squamata: Xenodermidae). Asian Herpetol Res, 12: 178–187.

Huelsenbeck J. P., Hillis D. M. 1993. Success of phylogenetic methods in the four-taxon case. Syst. Biol, 42: 247–264.

Li C, Yuan Z, Li H, et al. The tenth member of stout newt (Amphibia: Salamandridae: *Pachytriton*): Description of a new species from Guangdong, southern China. Zootaxa, 2018, 4399: 207–219.

Luedtke J. A., Chanson J., Neam K., Hobin L., Maciel A. O., Catenazzi A., Borzée A., Hamidy A., Aowphol A., Jean A., Sosa-Bartuano Á., Fong G. A., de Silva A., Fouquet A., Angulo A., Kidov A. A., Muñoz Saravia A., Diesmos A. C., Tominaga A., Shrestha B., Gratwicke B., Tjaturadi B., Martínez Rivera C. C., Vásquez Almazán C. R., Señaris C., Chandramouli S. R., Strüssmann C., Cortez Fernández C. F., Azat C., Hoskin C. J., Hilton-Taylor C., Whyte D. L., Gower D. J., Olson D. H., Cisneros-Heredia D. F., Santana D. J., Nagombi E., Najafi-Majd E., Quah E. S. H., Bolaños F., Xie F., Brusquetti F., Álvarez F. S., Andreone F., Glaw F., Castañeda F. E., Kraus F., Parra-Olea G., Chaves G., Medina-Rangel G. F., González-Durán G., Ortega-Andrade H. M., Machado I. F., Das I., Dias I. R., Urbina-Cardona J. N., Crnobrnja-Isailović J., Yang J. H., Jiang J. P., Wangyal J. T., Rowley J. J. L., Measey J., Vasudevan K., Chan K. O., Gururaja K. V., Ovaska K., Warr L. C., Canseco-Márquez L., Toledo L. F., Díaz L. M., Khan M. M. H., Meegaskumbura M., Acevedo M. E., Napoli M. F., Ponce M. A., Vaira M., Lampo M., Yánez-Muñoz M. H., Scherz M. D., Rödel M. O., Matsui M., Fildor M., Kusrini M. D., Ahmed M. F., Rais M., Kouamé N. G., García N., Legrand Gonwouo N., Burrowes P. A., Imbun P. Y., Wagner P., Kok P. J. R., Joglar R. L., Auguste R. J., Albuquerque Brandão R., Ibáñez R., von May R., Hedges S. B., Biju S. D., Ganesh S. R., Wren S., Das S., Flechas S. V., Ashpole S. L., Robleto-Hernández S. J., Loader S. P., Incháustegui S. J., Garg S., Phimmachak S., Richards S. J., Slimani T., Osborne-Naikatini T., Abreu-Jardim T. P. F., Condez T. H., De Carvalho T. R., Cutajar T. P., Pierson T. W., Nguyen T. Q., Kaya U., Yuan Z. Y., Long B., Langhammer P., Stuart S. N. 2023. Ongoing declines for the world’s amphibians in the face of emerging threats. Nature, 622: 308–314.

Lyu Z. T., Chen Y., Yang J. H., Zeng Z. C., Wang J., Zhao J., Wan H., Pang H., Wang Y. Y. 2020. A new species of Nidirana from the *N. pleuraden* group (Anura, Ranidae) from western Yunnan, China. Zootaxa, 4861: 43–62.

Lyu Z. T., Wang J., Zeng Z. C., Zhou J. J., Qi S., Wan H., Li Y. Y., Wang Y. Y. 2021. A new species of the genus *Tylototriton* (Caudata, Salamandridae) from Guangdong, southern China, with discussion on the subgenera and species groups within the genus. Vertebr Zool, 71: 697–710.

Nishikawa K., Jiang J. P., Matsui M. 2011b. Two new species of *Pachytriton* from Anhui and Guangxi, China (Amphibia: Urodela: Salamandridae). Curr Herpetol, 30: 15–31.

Nishikawa K., Jiang J. P., Matsui M., Mo Y. M. 2011a. Unmasking *Pachytriton labiatus* (Amphibia: Urodela: Salamandridae), with description of a new species of Pachytriton from Guangxi, China. Zool Sci, 28: 453–461.

Nishikawa K., Matsui M., Jiang J. P. 2012. A new species of *Pachytriton* from China (Amphibia: Urodela: Salamandridae). Curr Herpetol, 31: 21–27.

Ren Y., Lü Y., Hu J., Yin L. 2021. Geodiversity underpins biodiversity but the relations can be complex: Implications from two biodiversity proxies. Glob Ecol Conserv, 31: e01830.

Shen Y. H., Shen D. W., Mo X. Y. 2008. A new species of salamander *Pachytriton archospotus* from Hunan Province, China (Amphibia, Salamandridae). Acta Zoologica Sinica, 54: 645–652.

Thompson J. D., Gibson T. J., Plewniak F., Jeanmougin F., Higgins D. G. 1997. The CLUSTAL_X windows interface: flexible strategies for multiple sequence alignment aided by quality analysis tools. Nucleic Acids Res, 25: 4876–4882.

Wang S.P., Zhu W., Fan L. Q., Li J. Q., Li Y. M. 2017. Amphibians testing negative for *Batrachochytrium dendrobatidis* and *Batrachochytrium salamandrivorans* on the Qinghai-Tibetan Plateau, China. Asian Herpetol Res, 8, 190–198.

Weisrock D. W., Papenfuss T. J., Macey J. R., Litvinchuk S. N., Polymeni R., Ugurtas I. H., Zhao E., Jowkar H., Larson A. 2006. Mol Phylogenet Evol, 41:368–383.

Weng D. M., Zhang L., Chen X. D., Shen G. C., Zhang H. W., Zhang F. G., Yu M. J. 2009. Species diversity of Fagus hayatae community in Qingliangfeng National Nature Reserve. Journal of Zhejiang Forestry Science and Technology, 29, 1–6.

Wu Y., Wang Y., Hanken J. 2012. New species of *Pachytriton* (Caudata: Salamandridae) from the Nanling mountain range, southeastern China. Zootaxa, 3388: 1–16.

Wu Y., Wang Y., Jiang K., Chen X., Hanken J. 2010. Homoplastic evolution of external coloration in Asian stout newts (*Pachytriton*) inferred from molecular phylogeny. Zool Scr, 39: 9– 22.

Wu Y., Wang Y., Jiang K., Hanken J. 2013. Significance of pre-Quaternary climate change for montane species diversity: insights from Asian salamanders (Salamandridae: *Pachytriton*). Mol Phylogenet Evol, 66: 380–390.

Wu Y.H., Murphy R.W. 2015. Concordant species delimitation from multiple independent evidence: A case study with the *Pachytriton brevipes* complex (Caudata: Salamandridae). Mol Phylogenet Evol, 92: 108–117.

Yuan Z. Y., Zhang B. L., Che J. 2016. A new species of the genus *Pachytriton* (Caudata: Salamandridae) from Hunan and Guangxi, southeastern China. Zootaxa, 4085: 219–232.

